# Multiscale mechanical models of DNA nanotubes: A theoretical basis for the design and tuning of DNA nanocarriers

**DOI:** 10.1101/2021.12.12.472306

**Authors:** Han-Lin Liu, Neng-Hui Zhang, Wei Lu

## Abstract

DNA nanostructures are one of potential candidates for drug carriers due to their good biocompatibility and non-specificity in vivo. A reliable prediction about mechanical properties of artificial DNA structures is desirable to improve the efficiency of DNA drug carriers, however there is only a handful of information on mechanical functionalities of DNA nanotubes (DNTs). This paper focuses on quantifying the multiscale correlations among DNT deformation, packaging conditions and surrounding factors to tune mechanical properties of DNTs. By combining WLC statistical mechanics model, Parsegian’s mesoscopic liquid crystal model and Euler’s continuum beam theory, we developed a multiscale DNA-frame model; then theoretically characterize the initial packed states of DNTs for the first time, and reveal the diversity mechanism in mechanical properties of DNTs induced by interchain interactions and initial packed states. Moreover, the study of parameters, such as packaging conditions and environmental factors, provides a potential control strategy for tuning mechanical properties of DNTs. These conclusions provide a theoretical basis for accurately controlling the property and deformation of DNT in various DNT dynamic devices, such as DNA nanocarriers.

## 1. Introduction

Viruses and cancer have become the most challenging health burden for human beings [1, 2]. Traditional drug nanocarriers, such as synthetic polymers [3, 4], carbon nanotubes (CNTs) [5, 6] and viral capsids [7] are able to transport various functional agents into cells, while they are often hampered by the immune system and drug resistance of organism due to their non-specificity and potential toxicity [8, 9]. To address these issues, researchers focus on the artificial DNA nanostructures with good biocompatibility and tunable mechanical property [10–12] as new types of carriers. These DNA nanocarriers can effectively circumvent the drug resistance and avoid the residue of non-degradable elements [13, 14]. For example, DNA nanotubes can be taken up by viral infection or living cancer cells [15]; it can also regulate the vascular smooth muscle cell autophagy [16], and control the delivery of cargo [17]. The ever-increasing application of DNA structures with user-defined shapes and properties requires the detailed knowledge about their complex mechanical properties [18, 19].

DNA nanotubes (DNTs), as a common building block for higher-order structures, can be achieved by tile-assembly or by the DNA origami approach. And the state that the multiple double-stranded DNA (dsDNA) arranges into multihelix bundles [18, 19] is called the packaging state. An effective prediction of DNT’s loading/packaging property and stability/rigidity under different physiological conditions would be desirable to support the design and tuning of clinically suitable drug carriers [1, 16]. For single dsDNA molecules, Bao indicated that there were at least three different deformation regimes before phase transition from B-form DNA to S-form DNA [20]. With low applied forces, the global end-to-end extension behavior of dsDNA could be interpreted by the classical models of entropic elasticity of polymer [20, 21]; with intermediate forces, the deformability of dsDNA needs to be quantified by the WLC model [21, 22]; and with high longitudinal forces, the force-extension relationship of dsDNA is described by the e-WLC model [23]. For bulk dsDNA solutions, Manning and Geggier et al. showed the ionic-strength and temperature dependency of dsDNA persistence length, respectively[24, 25]; Parsegian et al. gave an effective interaction potential to characterize complex micro-interactions in dsDNA nematic liquid crystals[26, 27]. Compared with the comprehensive knowledge of single dsDNA molecules and bulk DNA solutions, there is only a handful of investigations on mechanical functionalities of DNTs.

Magnetic/optical tweezers experiments show a considerably increased bending rigidity of DNTs compared to a single dsDNA [19]. Simple predictions on the bending rigidities of DNTs [28–31], which were obtained from a simple mechanical model in the frame of continuum mechanics, were over-/under-estimated in certain conditions [30]. Kauert and Joshi et al. evaluated elastic properties of DNTs by Monte Carlo (MC) [19] and molecular dynamics (MD) [32] simulations, respectively, while the computational cost is relatively expensive. Moreover, experiments also reveal a strong enhancement in bending rigidity of DNTs compared with a small enhancement in torsional rigidity with the increase of helix numbers [19]. Although the finite element method (FEM) reproduced this discrepancy, the mechanism still remains an open question [19]. Ma et al. [33] measured rigidities of DNTs by AFM nano-indentation and ascribed this discrepancy to its anisotropic mechanical properties. Pfitzner et al. [34] predicted the modulus of DNTs based on the superposition principle, found that there was an order of magnitude bigger than the experimental values. This suggested the unfitness of WLC model in characterizing the behavior of multihelix bundles.

Besides the above-mentioned effect, other adjustable factors are also causing concern, such as DNT diameter [35], cation concentration and temperature [36]. However, the insufficient information on mechanical rigidity and stability of DNT hinders the rational design of DNA nanomaterials for applications. In the previous theoretical works, the interchain interactions between DNA cylinders were often neglected even for tightly packaged lattices [10]. Furthermore, the initial packaging state closely relevant to these microscale interactions was not well characterized, and the WLC model was directly applied without noticing the fact that multihelix DNA bundles are a sort of new artificial materials between single dsDNA molecules and bulk DNA solutions [20, 21].

In this paper, we aim to build a theoretical basis for the design of DNT mechanical properties via a prestressed tensegrity. First, DNT in magnetic tweezers experiments is molded as a pretensioned frame structure. Its initial packaging state is determined by a multiscale model considering energy contributions both from elastic deformation of a DNA thin-rod in WLC model [20, 21] and interchain interaction in the mesoscopic DNA liquid crystal model [27, 37]. Second, taking this initial packaging state as a benchmark, the force-extension curves of DNTs under longitudinal force and the stretch rigidities affected by the packaging conditions and environmental factors on stretch rigidity are explored by the principle of minimum potential energy. Then, the bending or torsional rigidities of DNTs are predicted analytically by using the theories for tension-bending-torsion combined deformations of DNA thin-rods, the force balance equations and deformation compatibility conditions at the end of DNA frame in pure bending or torsional state. The robustness [18] or sensitivity of bending/ torsional rigidities to the packaging and surrounding conditions are also checked, especially in response to the debate on the effect of helix numbers when interpreting the relevant experimental findings [19, 33].

## 2. Materials and methods

Magnetic tweezers [19], optical tweezers [34], fluorescence microscopy [18, 31] and atomic force microscope (AFM) [33] experiments are often used to measure the mechanical properties of DNTs. The description of these experimental methods here is valuable for the establishment of a new DNT model. Due to the limited space, first,we just introduce the magnetic tweezers experiment among these interesting works. Note that the experimental methods in the following sections are used from previous studies. Then, by collecting and fitting the experimental data of Schiffels [18], Kauert [19], Wang [31], Ma [33] and Pfitzner [34] et al., the stretching, bending and torsional rigidities of DNTs are given and analyzed. Finally, a multiscale model of DNTs is presented to reveal the diversity mechanism in mechanical properties of DNTs.

### 2.1 Experiments of DNT mechanical properties

#### 2.1.1 Materials and structures in magnetic tweezers experiment

Thanking for the good biocompatibility and the elegant principle of complementary base pairing of dsDNA, researchers can fabricate dsDNA into programmable DNTs with different regular polygon packaged patterns, such as 6-helix, 8-helix and 10-helix DNTs in Fig. 1a by the DNA origami or by tile-assembly approach.

**Fig. 1.**
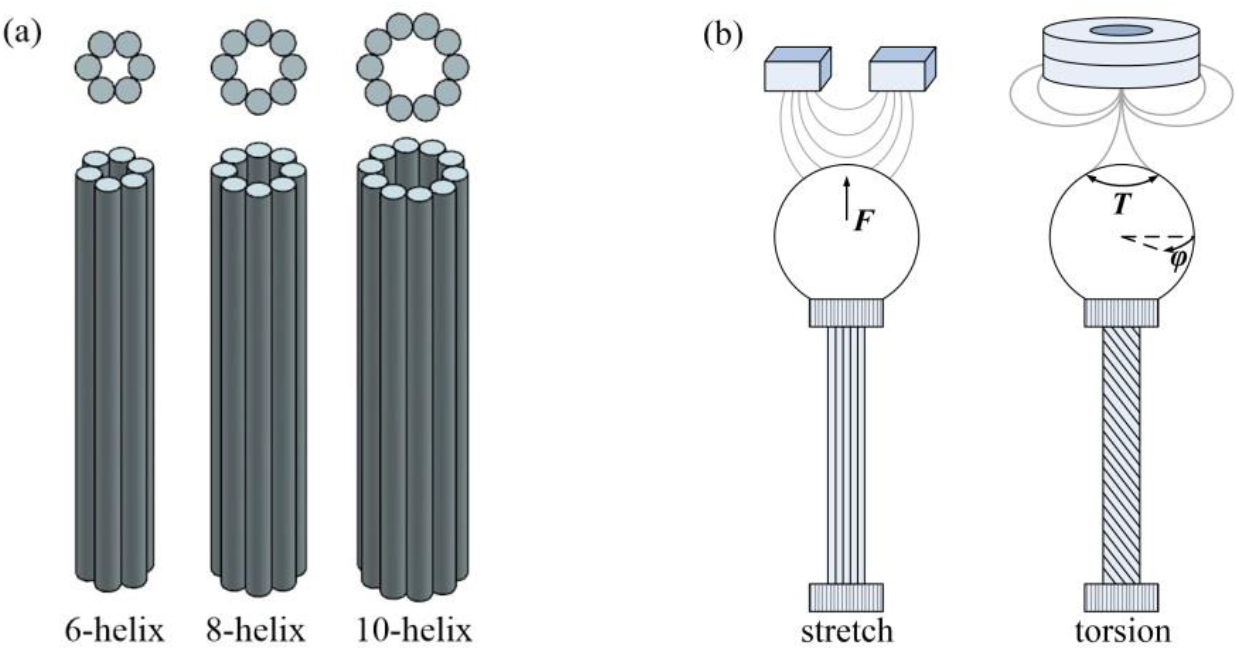
Schematic of (a) 6-helix, 8-helix, 10-helix DNTs and (b) DNT magnetic tweezers experiment

#### 2.1.2 Steps of magnetic tweezers experiment

As shown in Fig. 1b, one end of DNTs was bound to magnetic bead and flushed into a flow cell where the second end was attached to glass substrate. The position of magnetic bead was adjusted to apply the force/torque to DNTs, and the long time trajectories (e.g. 273 s) at a fixed force for 3D positions of the bead were tracked. Then, the force-extension or torque-rotation curves were recorded.

#### 2.1.3 Data analysis

We first collected the force-extension and torque-rotation curves in the previous experiments [18, 19, 31, 33, 34], then gave the rigidities of DNTs in Fig. 2 by fitting these curves.

**Fig. 2.**
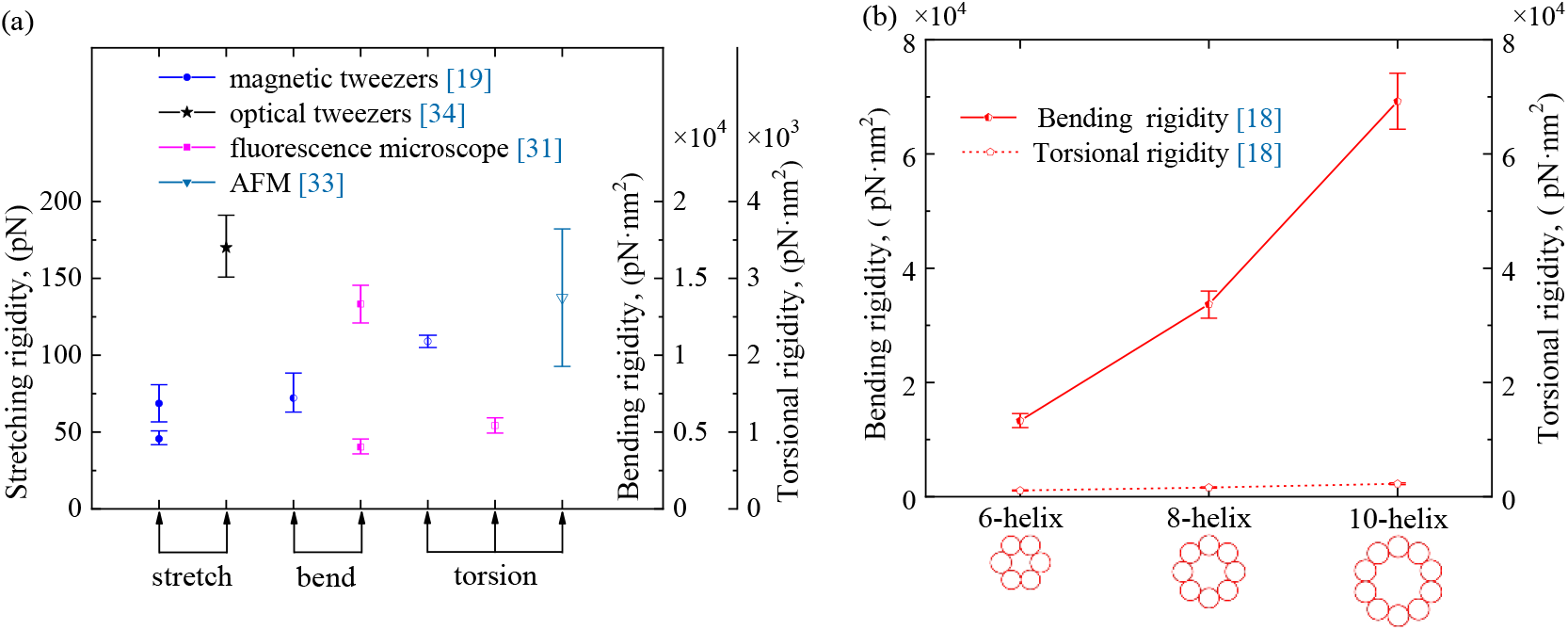
(a) Elastic rigidities of 6-helix DNTs under different experiments and (b) bending/torsion rigidities of DNTs with different helix number in fluorescence microscopy experiment. And the relevant environmental conditions and material information were obtained from the previous corresponding studies [18, 19, 31, 33, 34].

As shown in Fig. 2a, the stretching rigidity of 6-helix DNTs in magnetic tweezers experiment fluctuates from 41 pN to 80 pN, while the rigidity in optical tweezers experiment fluctuates from 150 pN to 191 pN. In addition, the bending rigidity and torsional rigidity of 6-helix DNTs in different experiments also have divergent fluctuation ranges. These diversities are ascribed to different environmental factors and their different structures.

As shown in Fig. 2b, in the fluorescence microscopy experiments, the bending and torsional rigidities of DNTs with different helix numbers have divergent fluctuation ranges. Furthermore, there exists a strong enhancement in bending rigidity while a small enhancement in torsional rigidity with the increase of helix mumbles. Obviously, the packaging conditions have a significant impact on DNT properties.

It can be seen that the tools, environmental factors and packaging conditions make the diversity of DNTs rigidities, while the experiments can only obtain the properties of DNTs under specific conditions. Here we will present a multiscale model of DNTs to characterize the influence of these factors on properties of DNTs.

### 2.2 Multiscale model of DNTs

#### 2.2.1 Simplified model for DNT magnetic tweezers experiment

Considering the architecture feature of DNTs, the constraint of substrate and magnetic bead in magnetic tweezers experiment, we model the DNT as a reduced frame structure with two rigid ends as shown in Fig. 3a [38], and each dsDNA chain is coarse-grained as a cylindrical rod with the radius of *r* and the packaging length of *L*_0_. We model the pre-stress as a longitudinal pre-stretching force *F*_s0_ applied at each dsDNA’s end [39], and the resultant of these pre-stretching forces creates the packaging force *F*_0_ of DNT. Furthermore, DNT carrier with user-designed shapes and tunable mechanical property can be obtained by designing its structural element with different packaging density (determined by interchain distance *d_i_*) [18] or geometrical pattern (determined by helix number *n* and each chain location). The local coordinate system of DNT was set up in Fig. 3b to analyze the complex deformation of each DNA rod when the external forces (tensile force *F*_s_, torque *T*_x_ and bending moment *M_z_*) are transferred from the rigid end to the cylinders.

**Fig.3.**
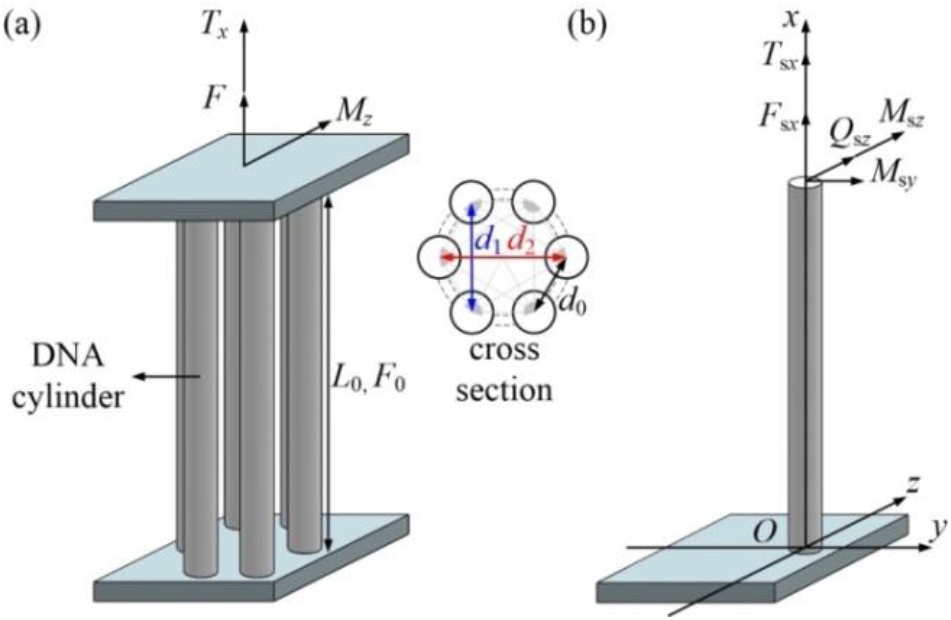
(a) Reduced frame structure of DNT in magnetic tweezers experiment and (b) local coordinate system for a single DNA cylindrical rod.

#### 2.2.2 Packaging state of DNTs

Based on the multiscale models of dsDNA molecules (see detail in Appendix A), a multiscale energy model of DNTs will be presented to characterize the packaging state of DNTs. The various constraints or forces applied during packaging process will lengthen the end-to-end distance of the molecule from only a few thousandths of the length of B-form dsDNA to its packaging length *L*_0_ [40]. In this transition process, the DNT’s energy variation, П_tb0_, includes the changes of DNA intrachain energy and interchain energy, i.e.

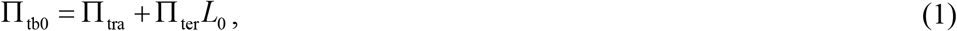

in which the change of intrachain energy П_tra_ can be indirectly characterized by [38]

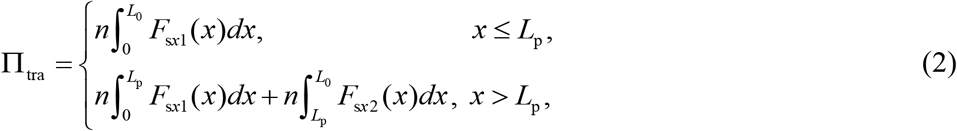

where *x* is the global extension with the same direction as the force *F*_s*x*_(*x*); the contour length *L* is the length when DNA chain is fully extended; the partition length *L*_p_ is the characteristic boundary between WLC and e-WLC models, and *L*_p_≈ 0.969*L*[40]; and the change of interchain energy per length, П_ter_, is described by Parsegian’s mesoscopic model in our previous studies [26, 37], 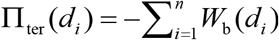.

Given the packaging interchain distance *d_i_*, the packaging length *L*_0_ could be determined by the minimum potential theory of energy. Then, substituting *L*_0_ into Eq. (A. 1) yields the effective longitudinal force *F_s0_* applied to each dsDNA. Summing all longitudinal forces of dsDNA yields the effective packaging force *F*_0_ of DNT as

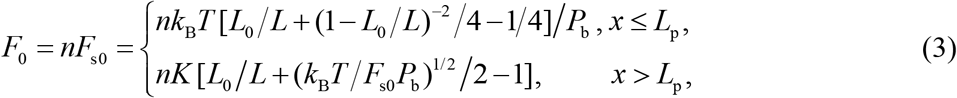

where *k_B_* is the Boltzmann constant, and *k*_B_=1. 38× 10^−23^ J/K, *T* the absolute temperature, *K* the stretch rigidity, and *K*= 4*P*_b_*k*_B_*T*/*r*^2^ [23]; the persistence length *P*_b_ defines the distance over which two segments of DNA chain remain directionally correlated. In addition, the critical load formula, *f*_cr_ = (2*P*_b_/*r*)^1/3^*k*_B_*T*/2*r* [23], is used to judge whether the deformation of single dsDNA chain is entropy or elasticity dominated.

#### 2.2.3 Tensile deformation of pretensioned DNT under longitudinal force

The global tensile deformation of single dsDNA displays different characteristics in response to different levels of longitudinal forces, so does the packaged DNT [41] induced by packaging conditions, as a particular assembly of multihelix dsDNA bundles. When DNT is subjected to external forces *F*_s_, it turns from its packaging state with length *L*_0_ to a new deformation state *L*_1_ and the total potential energy change is

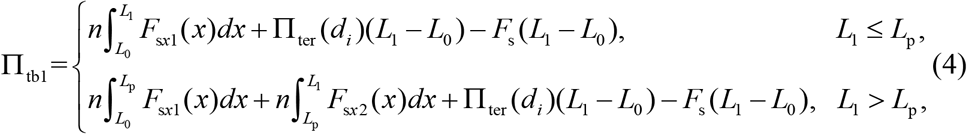

where *F*_s_(*L*_1_–*L*_0_) is the work of external force. By using the minimum potential energy principle, the extension of DNTs could be determined.

#### 2.2.4 Pure torsional state of pretensioned DNT

For a pure torsional state, i.e. *F* = 0, *M_z_* = 0 and *T_x_* ≠ 0 [19], determining the global deformation of each DNA rod is a statically indeterminate problem. As shown in Fig. 3b, the deformation compatibility condition at the top end is given as

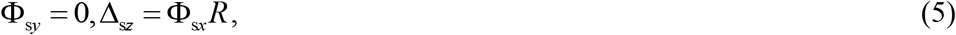

in which the radius of DNT is *R*= *d*_2_/2. When multihelix number *n* is even, the force and moment balance conditions along the *x*-axis for the rigid end are given as follows

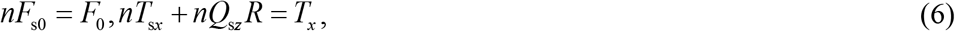

By combining Eqs. (5), (6) and Eqs. (B. 1), (B. 7), (B. 8), the end torsional angle of dsDNA rod can be given as

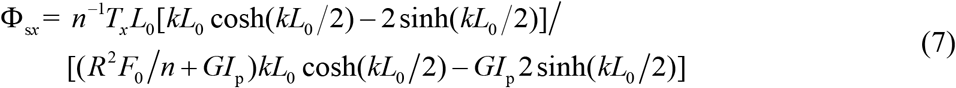

Given the hypothesis of rigid end, the end torsional angle of DNT equals to that of dsDNA rod, i.e. Φ_*x*_ =Φ_s*x*_, so the torsional rigidity of the pretensioned DNT is given as

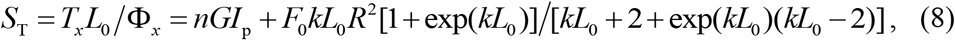

where the first term is contributed from the superimposing effect, and the second term is contributed from the tension-torsion-bending coupling effect related to the packaging state. In the case of neglecting pretension force, i.e. *F*_0_ ≈ 0, we get a simplified form of Eq. (11) as *S*_T0_ = *nGI*_p_ [1+6(*E*/*G*) (*R*/*L*_0_)^2^] (Eq. (C. 6), see more details in Appendix C).

#### 2.2.5 Pure bending state of pretensioned DNT

For a pure bending state, i.e. *F* = 0, *M_z_* ≠ 0 and *T_x_* = 0, the global deformation of each DNA rod could be easily obtained without involving the complex statically indeterminate problem. Under isomorphic hypothesis, as shown in Fig. 3b, from the moment balance along the *z*-axis, the relation between DNA rod’s end moment *M*_s*z*_ and the DNT’s external moment *M*_z_ is given as *M*_s*z*_ = *M*_z_/*n*. By using Eq. (5) and (B. 14), the curvature radius of each dsDNA rod at its local coordinate system is given as

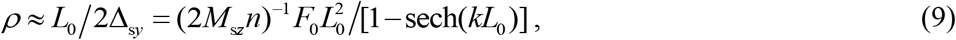

By combing Eq. (9) with *M*_s*z*_ = *M*_z_/*n*, the bending rigidity of each pretensioned DNA rod at its local coordinate system is denoted by

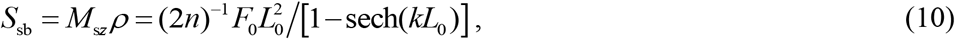

Then by using the parallel axis theorem and Eq. (10), the bending rigidity of the pretensioned DNT as to its centroidal principle axis is obtained as

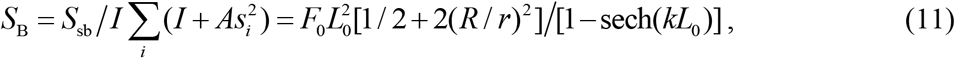

where *s_i_* is the distance from the mass center of each DNA rod’s cross-section as to the centroidal principle axis of DNT, the first term is contributed from the superimposing effect, and the second term is contributed from the eccentric effect. In the case of *F*_0_ ≈ 0, Schiffels et al. [18] gave a simple prediction, *P*_b_^*^ = *nP*_b_[1+2(*R*/*r*)^2^], for the persistence length of DNTs. Combining it with Eq. (A. 3), we get a degenerated form of Eq. (11)

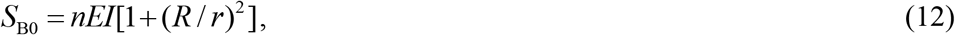

## 3. Results

### 3.1 The packaging performance of DNT

The robustness of packaging property to surrounding condition is investigated by using Eq. (3) and Eqs. (A. 1), (A. 2). As shown in Fig. 4, the packaging property curves show nonlinear variations, and they can be divided into entropy (I), WLC (II) and e-WLC (III) regimes according the deformation state of a single DNA rod in DNT homogenous deformation state based on critical value *f*_cr_ and partition length *L*_p_.

**Fig. 4.**
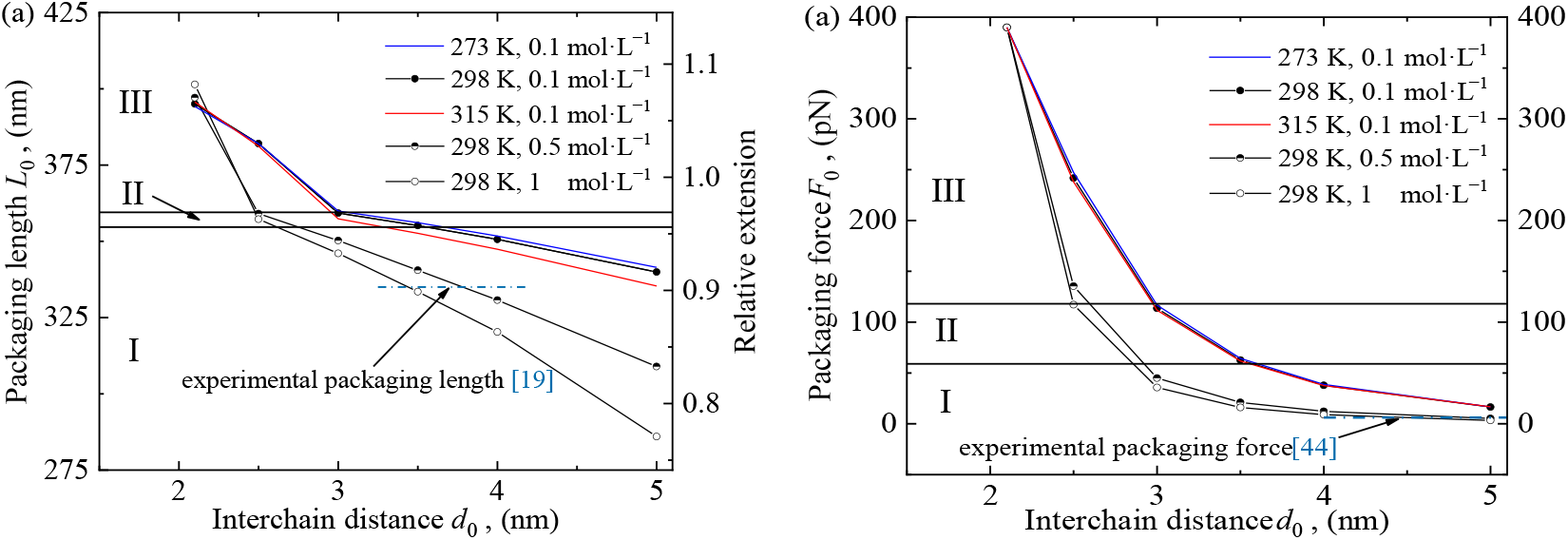
Packaging properties of DNTs under different interchain distances: (a) packaging length; (b) packaging force. And I, II and represent entropy, WLC and e-WLC regimes, respectively.

### 3.2 The elastic performance of DNT

The force-extension behavior of the packaged DNTs will be thoroughly studied by using Eq. (4) and Eqs. (A. 1), (A. 2). In calculation, DNT stretch rigidities are obtained by extracting the curve slopes of every three adjacent discrete points in these curves.

#### 3.2.1 Tensile performance

As shown in Fig. 5, with the increase of longitudinal force, the force-extension curve of DNT elevates its slope globally, which demonstrates a hardening process usually seen in biomaterials. The level of force, *F*_s_, decides the change of hardening rate with a relatively slow increasing trend in entropy (I) regime, a more dramatic one as approaching to the partition length *L*_p_ in WLC (II) regime, and a relatively stable, fast one after entering into e-WLC (III) regime.

**Fig. 5.**
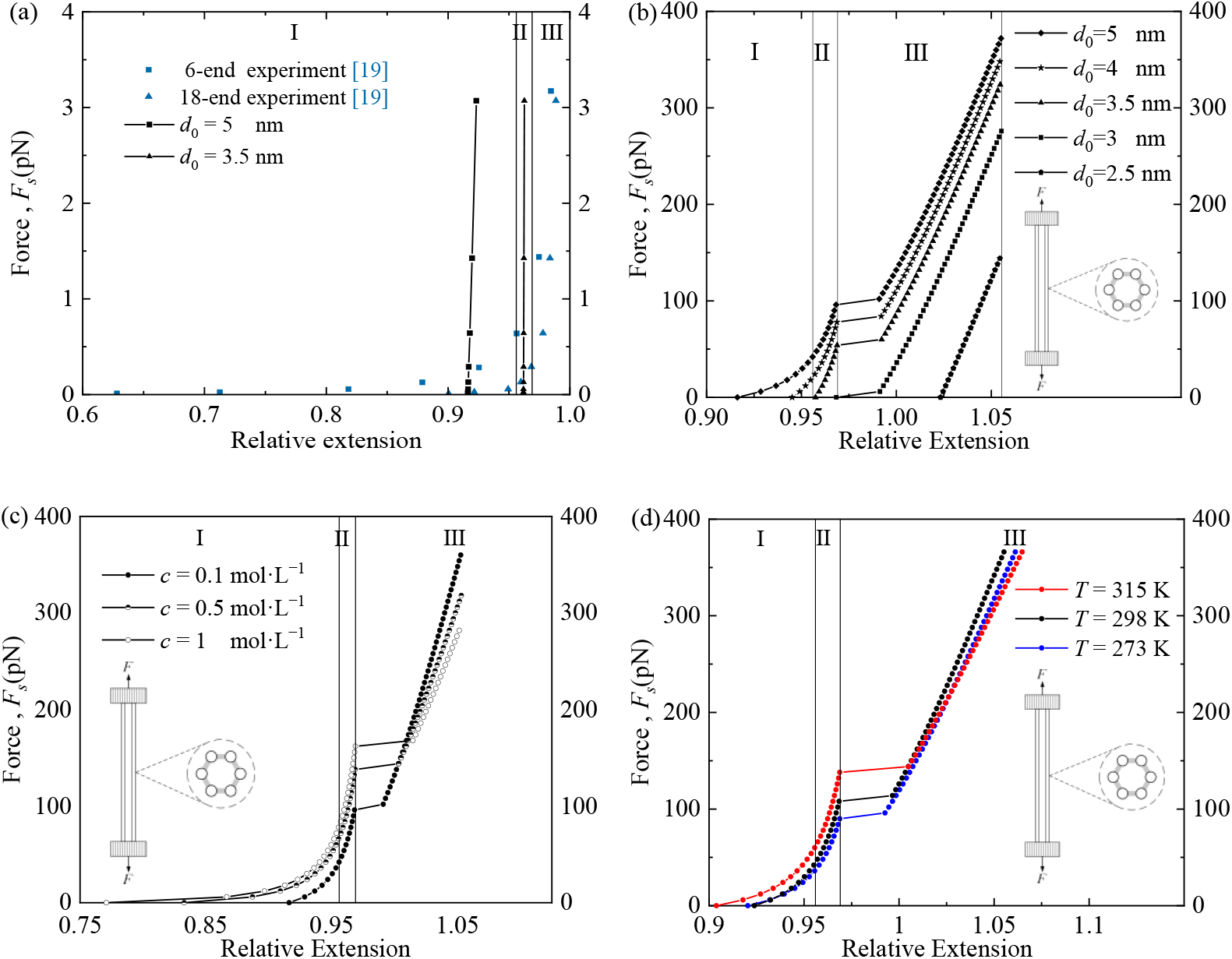
6-helix DNT force-extension curve (a) model predictions in comparison with experiment results [19]; model predictions with different (b)interchain distances (*T* = 298 K, *c* = 0.1 mol·L^−1^), (c)salt solutions (*T* = 298 K, *d*_0_ = 5 nm), (d) temperature conditions (*d*_0_ = 5 nm, *c* = 0.1 mol·L^−1^); and I, II and III represent entropy, WLC and e-WLC regimes, respectively. And the platform interval is caused by the original force gap between WLC model and e-WLC model for signal dsDNA chain.

The correlation between stretch rigidities of DNTs, initial packaging conditions and environment factors is shown in Fig. 6. A stronger external force will make DNT turning from entropy (I) regime to bending elasticity dominated WLC (II) regime with high stretch rigidity, and finally getting into stretch elasticity dominated e-WLC (III) regime with moderate stretch rigidity due to stiffness soften effect often seen in biomaterials and biomimetic materials [42, 43]. In addition, the larger packaging force may make DNT enter the WLC (II) or e-WLC (III) regimes directly.

**Fig. 6.**
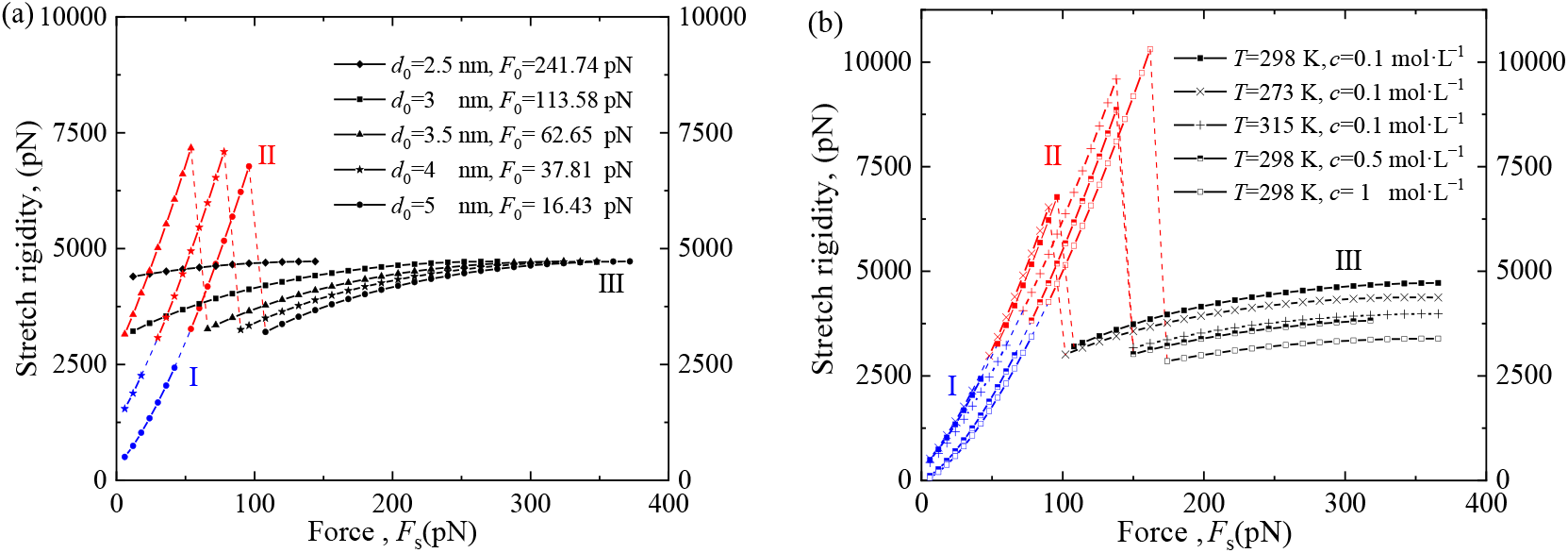
Variation of 6-helix DNT stretch rigidity with different (a) packaging conditions (*T*= 298 K, *c*= 0.1 mol·L^−1^) and (b) environmental factors (*d*_0_= 5 nm, *F*_0_= 16.43 pN) and external loads; and the blue, red and black lines represent entropy (I), WLC (II) and e-WLC (III) regimes, respectively. And the sudden drop between WLC regime and e-WLC regime is induced by the original force mismatch of WLC model and e-WLC model.

#### 3.2.2 Torsional and bending performance

The effects of packaging state on torsional/bending rigidity of DNT are shown in Fig. 7a. The variation tendency of torsional/bending rigidity versus packaging force is monotonic. Similarly to Fig. 4, the variation tendency of bending or torsional rigidity versus interchain distance in Fig. 7b decreases markedly at first, then turns to a slow state in an infection points (*d*_0_ =3.5 nm). In addition, it can be seen that the bending or torsional rigidity of DNTs decreases with the increase of salt concentration, and changes non-monotonously with the increase of temperature.

**Fig. 7.**
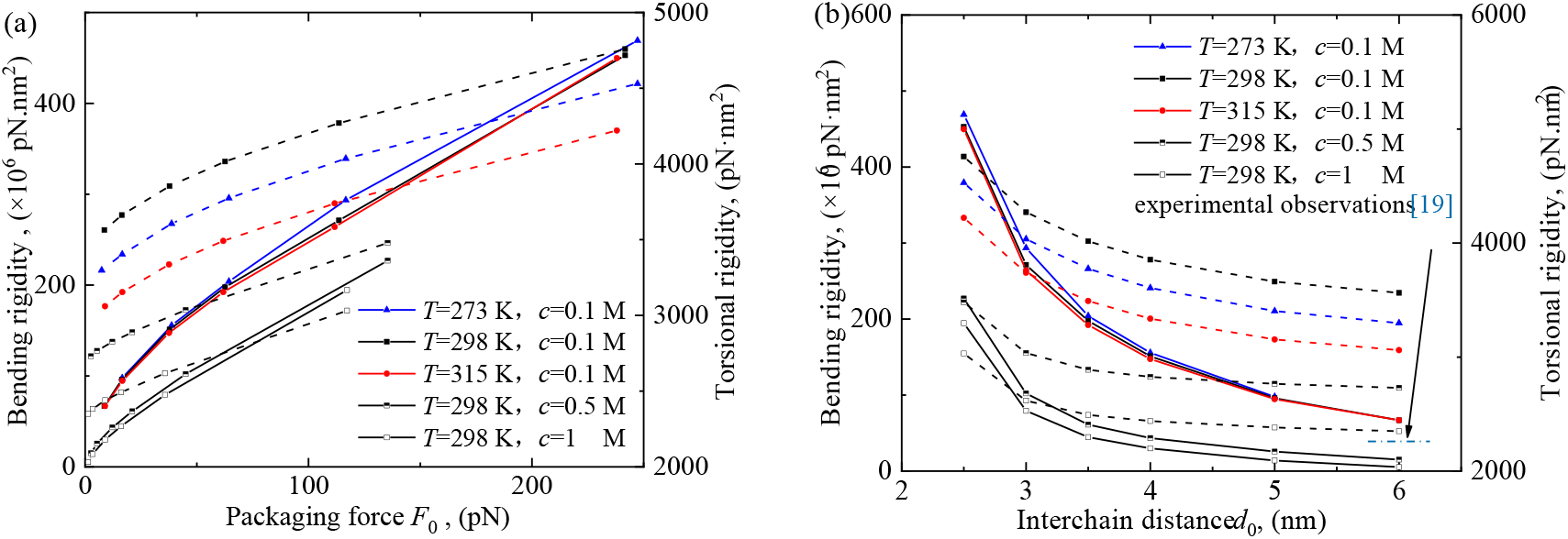
Variations of elastic rigidities of 6-helix DNT with (a) packaging force and (b) interchain distance; and the solid line represents bending rigidity, and the dotted line represents torsional rigidity.

### 3.3 The effect of helix number on rigidities of DNTs

Since it is difficult to present an analytic prediction of stretch rigidity of DNTs, the stretch rigidity considering packaging force is still obtained by the numerical fitting method as in Section 3.2, after the packaging states are determined; while the rigidity neglecting packaging force is obtained by multiplying the experimental data of single dsDNA [20] by helix number. As shown in Fig. 8, the stretch/bending rigidities can be strengthened by increasing helix number; while torsional rigidities are orders of magnitude smaller than bending rigidities, which endows the insensitivity of DNT torsional rigidity to helix number as found in previous experiments [18, 19, 31].

**Fig. 8.**
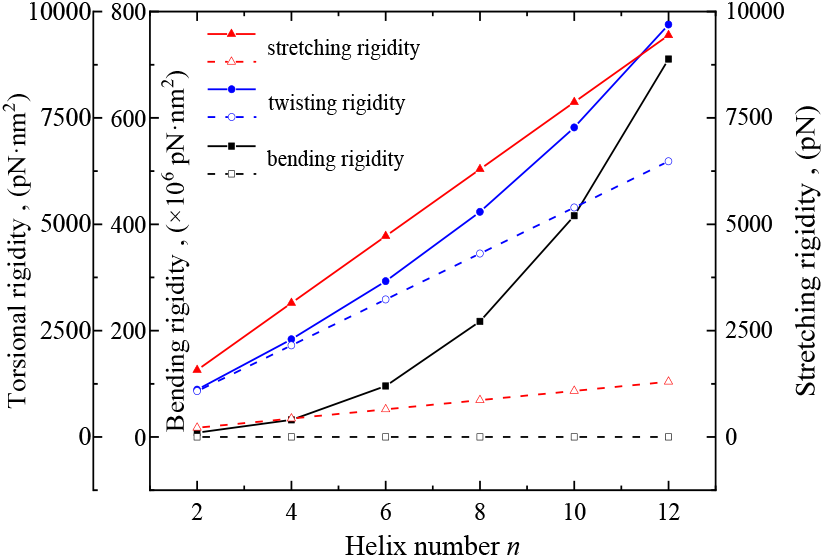
Variations of DNT elastic rigidities with helix number (*d*_0_= 5 nm, *c*= 0.1 mol·L^−1^, *T*= 298 K); and the solid and dotted line represent the predictions considering and neglecting packaging force effect, respectively.

## 4. Discussion

As for the packaging performance of DNT in Section 3.1 Fig. 4, our predictions approximate to the experimental result of 335 nm DNT [19] and the simple calculation from the experimental result of a single dsDNA [44]. The curves in Fig.4 suddenly slow their downward trends at the intersection between e-WLC (III) regime and WLC (II) regime. Meanwhile, the effect of concentration increase on packaging length also changes from promoting to inhibiting, and this inhibition becomes more obvious with the increase of interchain distance. However, the packaging properties are not sensitive to temperature changes. To reveal microscale mechanism in the above non-monotonic variation of DNT packaging property, the contribution of interchain interaction energy on DNT packaging properties in different deformation regimes is analyzed, and the mesoscopic energy ratio between intrachain energy and interchain energy are shown in Fig. 9a. Obviously the interchain interaction energy dominates the initial packaging states. The mesoscopic electrostatic energy and hydration energy jointly dominate in e-WLC (I) regime for tightly packaged patterns (*d*_0_< 3 nm), while only the electrostatic force dominates in WLC (II) regime when 3≤*d*_0_≤ 3.5 nm, then the conformational entropy takes over in entropy (I) regime for loosely packaged patterns (*d*_0_> 3.5 nm).

**Fig. 9.**
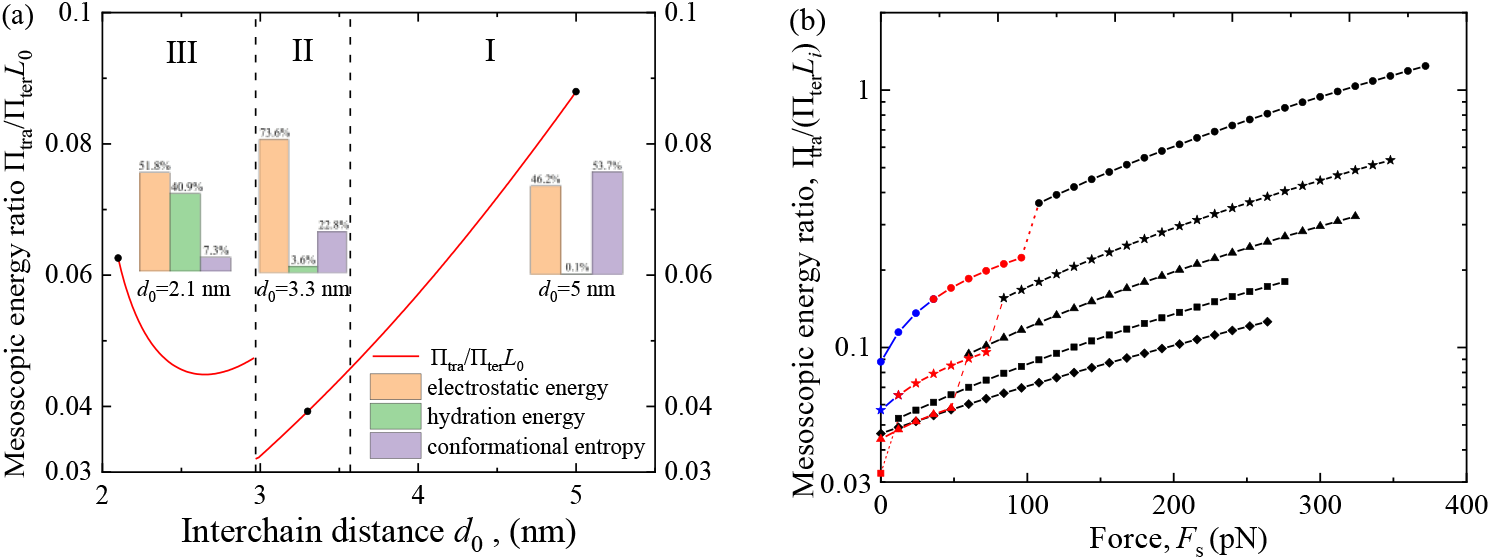
Mesoscopic energy ratio of DNT (a) in different deformation regimes and (b) under different external force when *c* =0.1 mol·L^−1^, *T* = 298 K; and I, II and III represent entropy, WLC and e-WLC regimes respectively; and the sudden change between II and III is induced by force mismatch of WLC model and e-WLC model.

As for the force-extension properties of DNTs in Section 3.2.1, our predictions of 6-helix DNT under low forces in Fig. 5a are close to the experimental results of DNTs under certain conditions [19]. And the packaging length, i.e. the initial extension of DNT will attain to above 90% of the contour length of single dsDNA. Base on the studies in Section 2.2.2, we deduce that the dominance transition of entropic, bending and stretch elasticity due to interchain interactions and packaging restrictions will make each DNA chain in DNT adopt more order and more straight conformation, different from the extension behavior of single dsDNA with the same length. Furthermore, the force-extension behaviors of DNTs under high forces are shown in Figs. 5b–5d for the first time. With the increase of initial packaging distance, salt concentration and absolute temperate, the extension gets smaller in entropy (I) and WLC (II) regimes. However, after entering into e-WLC(III) regime, it presents a monotonically increasing trend (Fig. 5c) or non-monotonous trend (Fig. 5d), respectively. These changes are the result of the competition between the decrease of interchain interactions (positively correlated) and the decrease of single dsDNA elastic rigidities (negatively correlated).

As for the quantitative analysis of DNT stretch rigidities in Section 3.2.1, our prediction under loosely packaging pattern (*d*_0_ > 3.5 nm) in Fig. 6a shows that the value of DNTs can attain to 2500 pN when *T* = 298 K, *c* = 0.1 mol·L^−1^, which is greater than the previous simple prediction of 250 pN obtained just by multiplying the experimental data of single dsDNA by multihelix number [45]. It suggests that the influence of interchain interactions on stretch rigidity cannot be ignored. In addition, the study of surrounding factors on stretch rigidities shown in Fig. 6b when *d*_0_=5 nm and *F*_0_ =16.43 pN reveals that the stretch rigidity decreases monotonously with the increase of salt concentration, whereas changes non-monotonously with the increase of temperature. It indicates that the decreasing interchain interaction can soften DNT with flexible tensile properties. Furthermore, the dominance transitions of entropic, bending and stretch elasticity, and the maximum stretch rigidity of DNTs (5000 pN) are consistent with the results given by Lin et al. [46] and the MD simulation results given by Joshi et al. [32]. To explain this phenomenon, the variation of mesoscopic energy ratio between intrachain energy and interchain energy under external loads is shown in Fig. 9b. Obviously, the intrachain interaction energy gradually dominates stretch rigidity of DNT that undergoes linear increase then stiffness softening. It indicates that the stretch rigidity of DNT is determined by the competition between the intrachain and interchain interactions, similar to the behavior found in biopolymer networks [47].

As for the DNT torsional rigidity in Section 3.2.2, our predictions agree well with the order of the experimental observations at room temperature [19] (Fig. 5b). From the second term of Eq. (8), we know that the monotonic variation of torsional/bending rigidity in Fig. 7a is due to the effect of packaging force. It has positive effect on torsional/bending rigidity firstly, and slows its effect gradually due to nonlinear coupling effect. The infection points (*d*_0_ = 3.5 nm) in Fig. 7b can be explained by using the same analytical methods as in Fig. 9. The emergence of infection points is because the electrostatic force exceeds hydration and conformational forces when *d*_0_≤3.5 nm, and the conformational entropy takes over the dominated regime when *d*_0_ >3.5 nm.

As for the elastic rigidities of DNTs in Section 3.3, all predictions neglecting the packaging force effect underestimate DNT elastic rigidities (Fig. 8). From the model in Section 2.2.4, we know that this is because the pretension-related packaging forces can improve stretch/torsional rigidities by several times while bending rigidities by thousand times. But the linear variation of stretch rigidity with helix number does not provide the rationality of superposition principle used in previous simple estimation due to existence of the pretension effect.

## 5. Conclusions

By providing multiscale DNA-frame models and characterization methods for accurately controlling deformation and property of DNT in various DNT dynamic devices under different clinically suitable environments, we compared DNT force-extension curves of magnetic tweezers experiments and theoretical predictions, and theoretically characterized packaging properties of DNT with different initial interchain distance or under different surrounding factors for the first time. DNA nanotubes are achieved by arranging multiple dsDNA into multihelix bundles, its loading capacity for nanomedicine is determined by interchain distance (Fig. 3a). However, influenced by the competition between interchain interaction and intrachain interaction of single dsDNA, the modulation of interchain distance will also change the robustness of DNTs. Although the increase in interchain distance can improve the drug loading capacity of DNT, it will also bring DNT to a loosely packaging pattern, in which the robustness to packaging condition should be paid attention especially.

Multiscale models of DNA nanotubes can be used not only to verify the existing experiments, but also to predict the situations that have not been experimented. For example, the predictions of DNT flexible tensile properties tuned by solution conditions reproduce well the trends found in the experiments of Kauert et al. [19], whereas those e-WLC curves at high longitudinal forces are displayed for the first time, which needs to be further validated by the related DNT experiments; the analyses of DNT torsional and bending rigidities also clarify the controversy on the dependency of rigidities on helix number [19, 33].

The multiscale model can not only provide a theoretical base for accurate control of nanomechanical deformation in various DNT dynamic devices, but also point out a new pathway to obtain optimal and robust elastic rigidities of DNTs. Take DNT in this article as an example, the competition in microscale interactions and nonlinear coupling effect of packaging force and pretensioned deformation make the interchain distance of 3.5 nm as an inflection point to low stretch rigidity, and also as the best interchain distance to improve torsional/bending rigidity when *c* = 0.1 mol·L^−1^.

However, there are still plenty of uncultivated areas worthy of being studied in the future, such as the shear effect on DNT torsional rigidity with the consideration of Holliday junctions, and accurate control of DNT critical buckling condition.

## Acknowledgments

This work was supported by the National Natural Science Foundation of China (Grant Nos. 11772182; 11272193; 10872121), and the Program of Shanghai Municipal Education Commission (Grant No. 2019-01-07-00-09-E00018).

## Appendix A Multiscale models of dsDNA molecules

For the tensile deformation of a single dsDNA molecule, its force-extension relationship is described by means of WLC [21, 22] and e-WLC models, i.e. [23]

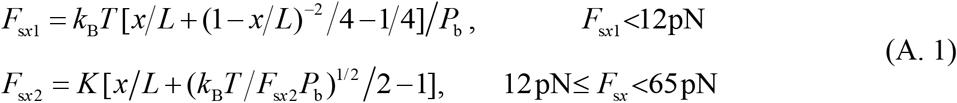

The equations introduced by Manning [24] and Geggier et al. [25] are used to consider ionic-strength dependency and temperature dependency of dsDNA persistence length *P*_b_ respectively, i.e.

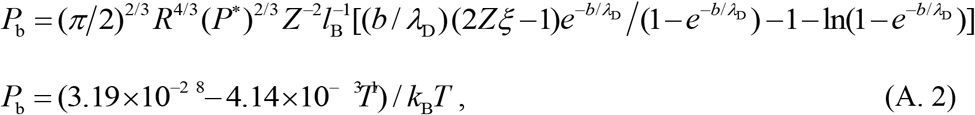

where *P*\* is the buckling persistence length, which is assumed to be insensitive to strength; *Z* is the cation valence; *ξ* is the dimensionless polyion charge density, and *ξ* = *l*_B_/*b*, where *b* the spacing between two charge sites, *l*_B_=*q*^2^/(4*πεk_B_T*), the Bjerrum length for the pure solvent, *q* the protonic charge, and *q*=1. 6×10^−19^ C, *ε* the dielectric constant of solvent and *ε*=*ε*_r_×*ε*_0_, *ε*_r_ the relative dielectric constant for water, and *ε*_r_ ≈ 80, *ε*_0_ the permittivity of vacuum, and *ε*_0_=8.85×10^−12^ F/m; *λ*_D_ is the Debye length, and 1/*λ*_D_=(*q*^2^ *β*∑*n_i_z_i_*/*ε*)^1/2^, where *β*=1/(*k*_B_*T*), *n_i_* the ionic concentration, *Z_i_* the ionic valence. A simple numerical formula for the Debye length in aqueous univalent salt solution of molarity *c* is given as *λ*_D_ =0.308 nm/(*c* mol·L^−1^)^1/2^.

To evaluate the torsional/bending rigidities of DNTs, we need know the global deformation of DNA rods. As shown in Fig. 3a, when the rigid end constraint is relieved and the corresponding support reaction forces/moments are applied at the top ends, each DNA rod is in a combined deformation, and has the same deformation under a given symmetrical packaged pattern with even helix numbers. We treat DNA rod as a homogeneous isotropic linear elastic body with Young’s modulus *E*, shear modulus *G*, cross-section area *A*= *πr*^2^, inertia moments *I*= *πr*^4^/4, polar moment of inertia *I*_p_= *πr*^4^/2. And the ability to resist deformation of dsDNA rod is characterized in terms of torsional rigidity *GI*_p_ and bending rigidity *EI*. The relationship between these rigidities and the persistence lengths is given by Odijk and Theo as follows [23]

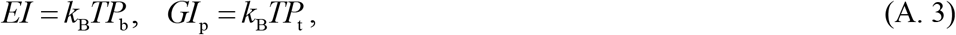

in which the torsional persistence length is taken as *P*_t_ ≈ 2.4*P*_b_ [18–20].

In the case of a tightly packaged DNT [10, 18], microscale interactions dominates local deformation of single dsDNA [48] and has the effect on global mechanical responses of DNTs [33]. Here we use the mesoscopic interaction potential energy per length *W*_b_ in terms of averaged interchain distance *d_i_* given by Parsegian et al.[26]

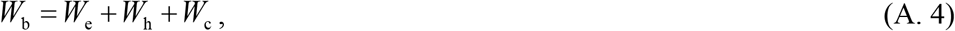

in which *W*_e_, *W*_h_, *W*_c_ represent, respectively, electrostatic energy, hydration energy, conformational entropy. The empirical equations and parameters for these energies of dsDNA liquid crystals in NaCl solutions can be obtained from previous works [26, 37].

## Appendix B Torsional and bending deformations of a cantilever DNA rod

As shown in Fig. 3b, subjected to the end external torque *T*_sx_, the cantilever DNA rod produces the torsional deformation along the *x*-axis, and the free-end rotation angle Φ_s*x*_ is given as

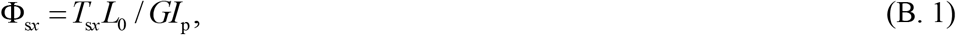

in which *GI*_p_ is the torsional rigidity of single DNA rod.

When the longitudinal packaging force *F*_s0_, the transverse external force *Q*_s*z*_ and the external moment *M*_s*y*_ are applied to the free-end, the cantilever DNA rod produces a bending deformation on the coordinate plane of *xoz*. Its deflection *w*_z_ is governed by

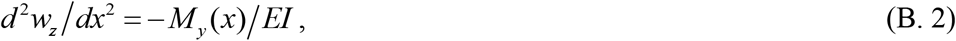

in which the internal moment *M_y_*(*x*) is given as

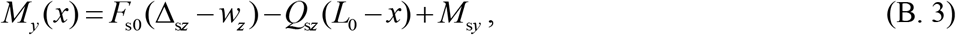

in which Δ_s*z*_ is the free-end deflection along the *z*-axis direction. Substituting Eq. (B. 3) into Eq. (B. 2) yields the following governing equation

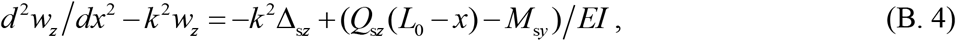

in which *k*^2^ = *F*_s0_/*EI*. The corresponding boundary conditions are given as

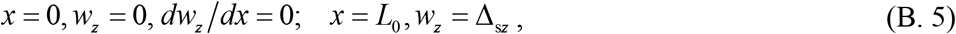

Combining Eq. (B. 4) and Eq. (B. 5), we can get

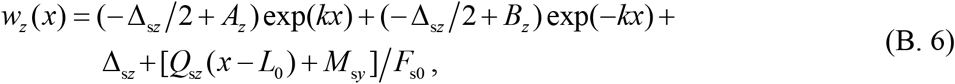

in which *A_z_* = [*Q*_s*z*_(*L*_0_*k*–1) –*M*_s*y*_*k*]/2*F*_s0_*k*, *B_z_*= [*Q*_s*z*_(*L*_0_*k* + 1) −*M*_s*y*_*k*]/2*F*_s0_*k*.

By using Eq. (B. 5) and Eq. (B. 6), we can obtain the free-end deflection as

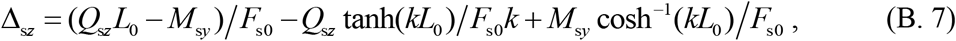

and the free-end rotation angle as

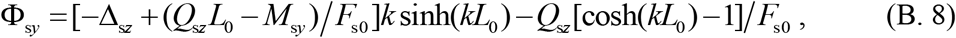

When the longitudinal packaging force *F*_s0_, and the external moment *M*_sz_ are applied to the free-end, the cantilever DNA rod produces a bending deformation on the coordinate plane of *xoy*, and its deflection *w_y_* is governed by

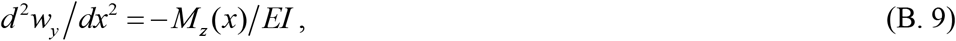

in which the internal moment *M_z_*(*x*) is given as

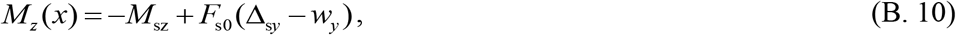

where Δ_s*y*_ is the free-end deflection along the *y*-axis direction. Substituting Eq. (B. 10) into Eq. (B. 9) yields the following governing equation

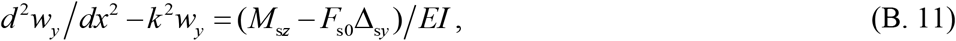

The corresponding boundary conditions are given as

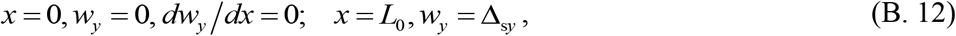

Combining Eq. (B. 11) and Eq. (B. 12), we can get

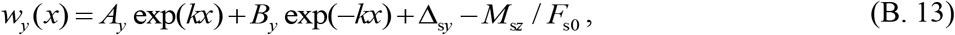

in which *A_y_* = *B_y_* =(−Δ_s*y*_ + *M*_s*z*_/*F*_s0_)/2.

By using Eq. (B. 12) and Eq. (B. 13), we can obtain the free-end deflection as

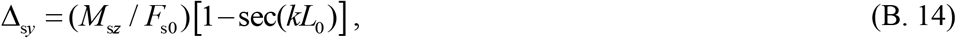

and the free-end rotation as

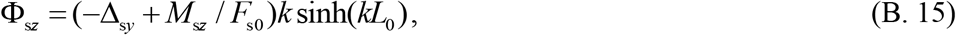

## Appendix C Torsional rigidity of pretensioned DNT neglecting pretension effect

In a pure torsional state, i.e. *F* = 0, *M_z_* = 0 and *T_x_* ≠ 0 [19], the end of each DNA rod is only subjected to transverse external force *Q*_s*z*_ and external moment *M*_s*y*_. If the packaging force *F*_s0_ could be neglected, and the cantilever DNA rod produces a bending deformation on the coordinate plane of *xoz* with its deflection *w_z_* still governed by Eq. (B. 2), however, the internal moment *M_y_*(*x*) is changed into

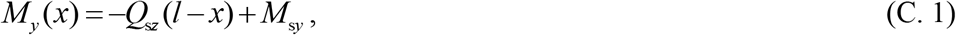

Combining Eqs. (B. 2), (C. 1) and the boundary conditions in Eq. (B. 5), we can obtain the free-end deflection Δ_s*z*_ and the free-end rotation angle Φ_*sy*_ as

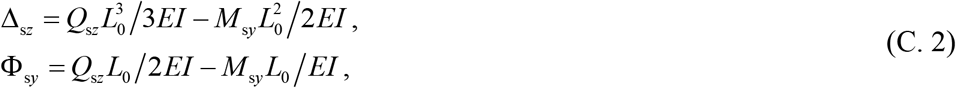

Combining Eqs. (B. 1), (C. 2) with Eqs. (6), (7), we obtain the following relations between the end forces/moments *T*_s*x*_, Q_s*z*_, *M*_s*y*_ and the external moment *T_x_*

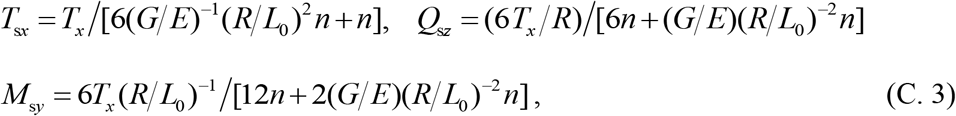

Substituting Eq. (C. 4) into Eq. (B. 1) yields the free-end torsional angle of DNA rod as

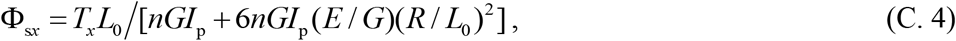

Given the rigid end hypothesis, the free-end torsional angle of DNT equals to that of DNA rod, i.e. Φ_*x*_ = Φ_s*x*_, so the torsional rigidity of pretensioned DNT is given as

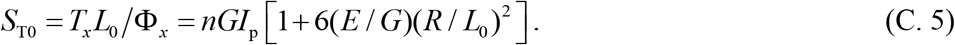

